# Widespread evolution of poricidal flowers: A striking example of morphological convergence across flowering plants

**DOI:** 10.1101/2024.02.28.582636

**Authors:** Avery Leigh Russell, Rosana Zenil-Ferguson, Stephen L. Buchmann, Diana D. Jolles, Ricardo Kriebel, Mario Vallejo-Marín

## Abstract

The evolution of tube-like floral morphologies that control pollen release via small openings (functionally poricidal flowers) represents a taxonomically and geographically widespread instance of repeated and independent evolution of a functionally similar morphology. Poricidal flowers are also often closely associated with buzz pollination by bees. Yet we lack an updated angiosperm-wide survey of their phylogenetic distribution. We identify all known angiosperm genera containing poricidal flowers via a literature survey. We determined their phylogenetic distribution and minimum number of independent gains and losses via a species-level angiosperm-wide phylogeny. We estimated if evolution of poricidal flowers is associated with changes in speciation/extinction via diversification rate analyses. Poricidal flowers occur across 87 angiosperm families and 639 genera containing >28,000 species. At the species level, an average of 205 independent gains and 215 losses of poricidal flowers occurred. Angiosperm-wide analyses suggest an early burst in poricidal evolution, but no differences in net diversification (origination-extinction) between non-poricidal and poricidal taxa. Analyses for two focal families however indicate strong context-dependent effects of poricidal flowers on diversification. Poricidal evolution thus represents a large-scale example of convergent evolution in floral form, but effects on diversification appear to be strongly contingent on phylogenetic and ecological background.

## Introduction

Patterns of convergent evolution are of fundamental interest in evolutionary biology. Broad convergence on functionally equivalent morphology across disparate groups is particularly common in plants, as in the case of succulent morphology or carnivorous pitcher plant traps, and has helped elucidate shared selective pressures, mechanisms, and developmental biases (Bennici, 2003; Ellison & Gotelli, 2009; Thorogood *et al*., 2018). A very geographically and taxonomically widespread pattern of convergent evolution is tubular floral morphology that restricts pollen dispersal via small openings, i.e., functionally poricidal floral morphology (hereafter ‘poricidal flowers’; Fig. 1) (Vallejo-Marin & Russell, 2024). Previous studies estimate that approximately 22,000 plant species possess poricidal anthers (‘poricidal species’), spread across 72 families and 544 genera (Buchmann, 1983). Additionally, flower traits that control pollen dispersal are key to angiosperm reproductive success (Hargreaves *et al*., 2009; Minnaar *et al*. 2019) and many poricidal plant species are also agriculturally important, such as tomatoes, cranberries, blueberries, and kiwifruit (Cooley & Vallejo-Marín, 2021). Yet the phylogenetic distribution and evolutionary patterns of poricidal flowers has not been formally characterized and the latest published taxonomic survey of poricidal flowers is now over 40 years old (Buchman, 1978; Buchmann, 1983).

**Figure 1.**
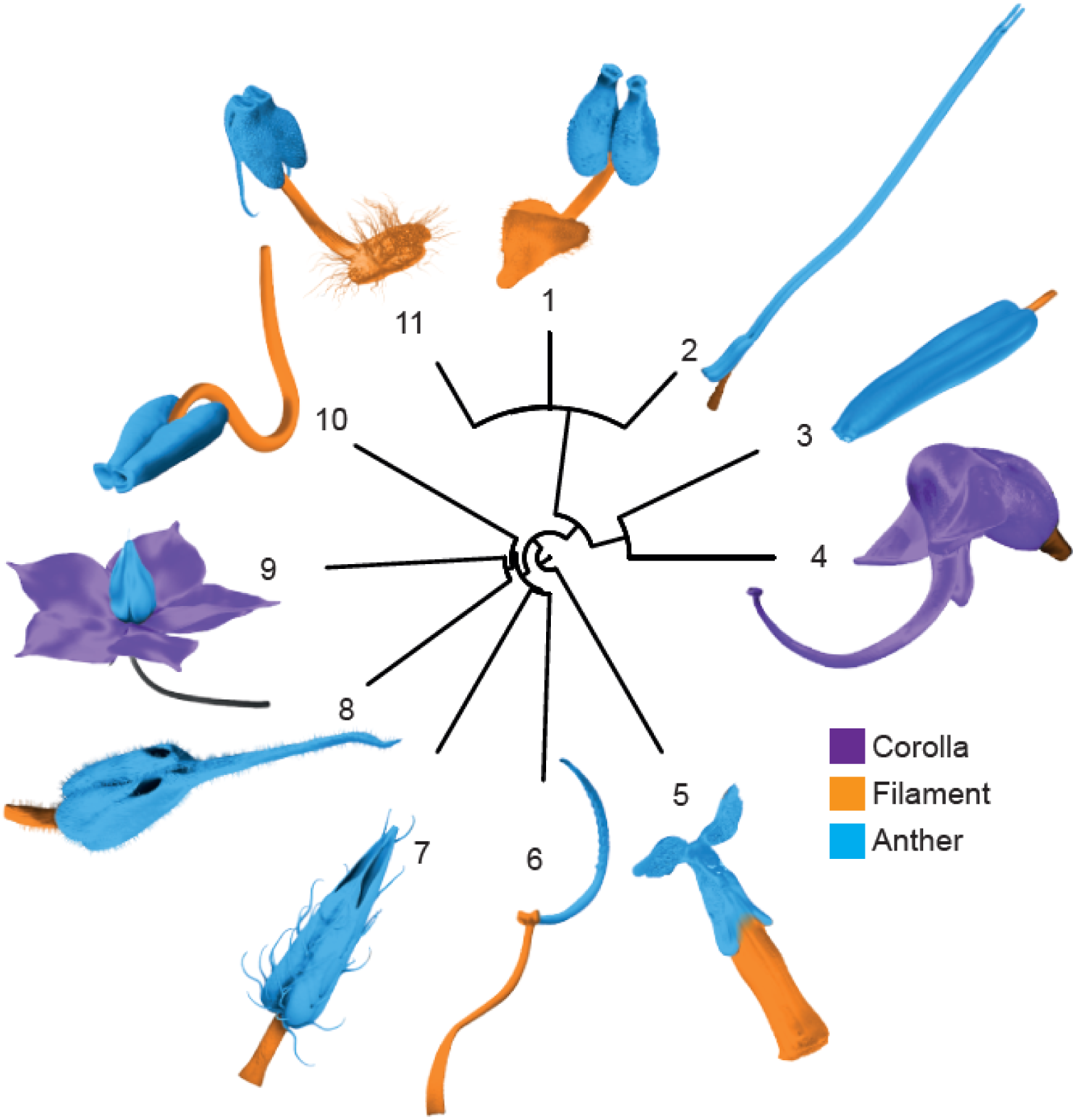
Diversity of poricidal flower structures illustrated across 11 taxa. The images depict stamens or whole flowers of taxa with poricidal flowers. For illustration of the diverse ways to build a poricidal flower, false colors indicate the structure or tissue involved: Corolla, Filament and accessory tissue (Filament), and Anther. The phylogeny at the center indicates relationships among taxa. Species as follows: 1 = *Chimaphila umbellata* (Ericaceae), 2 = *Demosthenesia cordifolia* (Ericaceae), 3 = *Solanum elaeagnifolium* (Solanaceae), 4 = *Pedicularis groenlandica* (Orobanchaceae), 5 = *Berberis sp*. (Berberidaceae), 6 = *Pleroma urvilleanum* (Melastomataceae), 7 = *Apeiba tibourbou* (Malvaceae), 8 = *Sloanea terniflora* (Elaeocarpaceae), 9 = *Sauvagesia capillaris* (Ochnaceae), 10 = *Senna* sp. (Fabaceae), 11 = *Arctostaphylos* sp. (Ericaceae). Redrawn from original 3D models by William Singleton and Daniel Hornung.

Poricidal flowers share a common functional morphology in which pollen is held within tubular floral structures with small openings (e.g., pores, valves, or slits; hereafter ‘pores’) through which the pollen is dispersed. In most cases, the tubular structure is composed of modified enlarged anthers, but may alternatively consist of modifications to the corolla or even the entire flower (Fig. 1) (Harris, 1905; Michener, 1962; Buchmann, 1983; De Luca & Vallejo-Marin, 2013; Russell *et al*., 2017; Dellinger *et al*., 2019b; Vallejo-Marin, 2019a). Poricidal flowers have been studied for over a century (Harris, 1905) and are frequently associated with a suite of floral characteristics (Buchmann, 1983; Renner, 1989; De Luca & Vallejo-Marin, 2013). Most poricidal plant species are animal pollinated, nectarless, and have dry, loose pollen (reduced pollenkitt lipids) and many are associated with enantiostyly (style handedness) and heteranthery (multiple kinds of stamens) (Buchmann, 1983; Vallejo-Marin *et al*., 2010). Furthermore, poricidal flowers are often closely associated with the buzz pollination interaction (although not all poricidal flowers are buzz pollinated; see Vallejo-Marin and Russell (2024) for some exceptions), in which pollination is nearly exclusively performed by bee species capable of rapidly contracting their indirect flight muscles while on the flower, generating strong vibrations that expel pollen onto the bee’s body (Macior, 1964; Buchmann, 1983; Proenca, 1992; Vallejo-Marin, 2019a; Vallejo-Marin & Russell, 2024). Indeed, poricidal flowers are thought to be a key pollen resource for these buzzing bees, particularly in tropical regions (Delgado *et al*., 2023). Poricidal floral morphologies may facilitate speciation by filtering out less effective pollinators (Cardinal *et al*., 2018). Yet because of specialization on a subset of pollinators, poricidal morphology may make these angiosperm taxa more vulnerable to extinction (Renner, 1989; Pacheco Filho *et al*., 2015; Dellinger *et al*., 2019a; Vallejo-Marin, 2019b). Consequently, poricidal floral morphologies are consistently suggested to be adaptations to pollinators and thus they might be key traits affecting patterns of speciation and extinction.

Previous attempts at comprehensively studying poricidal flowers were restricted to species with pored anthers presumed to be buzz pollinated (Harris, 1905; Buchmann, 1983). However, functionally poricidal floral morphologies are more diverse (Fig. 1). Additionally, clades of poricidal species can comprise thousands of species (e.g., *Solanum*, Melastomataceae) and the rate of gain and loss of poricidal floral morphologies has yet to be quantified. Although poricidal species are extremely widespread, clades are often all poricidal or not and thus studies of small phylogenetic clades suffer from the limitations of rarely seeing evolutionary transitions to and from poricidal flowers. Consequently, large-scale analyses are needed to allow generalization and provide sufficient statistical replication to test the effect of poricidal flower morphologies on diversification (Beaulieu & O’Meara, 2018; ZenilLFerguson *et al*., 2019; Joly & Schoen, 2021). However, large-scale phylogenetic analyses bring their own challenges (Helmstetter *et al*., 2022). First, it is challenging to obtain information on multiple traits for thousands of species (i.e., presence or absence of poricidal flowers). Second, well resolved mega-phylogenies are required. Fortunately, recent angiosperm-wide phylogenies have become available (Sauquet & Magallón, 2018; Jin & Qian, 2019; Janssens *et al*., 2020). Third, running complex diversification models in large phylogenies is computationally demanding and proven methods continue to be developed to speed up large-scale and complex analyses. Finally, the effects of a single trait in a large tree can be confounded with heterogeneity in the process of diversification by other unmeasured factors (Beaulieu & O’Meara, 2018; Beaulieu & O’Meara, 2019; Donoghue & Edwards, 2019).

Here, we investigate the evolutionary history of poricidal flowers across angiosperms to address the following questions. **1)** What proportion of angiosperms possesses poricidal flowers? **2)** How many times have poricidal flowers been gained and lost over the evolutionary history of flowering plants? **3)** Is the presence of poricidal flowers associated with changes in net diversification (speciation minus extinction) and diversification rates across all angiosperms? And **4)** Is the presence of poricidal flowers associated with diversification rates (speciation-extinction) within some key families? We therefore conducted an extensive literature survey to identify all presently known angiosperm genera containing species with poricidal flowers. We then used a species-level angiosperm-wide phylogeny to determine the phylogenetic distribution of poricidal flowers, the geologic ages of poricidal lineages, and the number of independent gains and losses of functionally poricidal floral morphology. We also conducted diversification rate analyses to estimate if the evolution of poricidal flowers is associated with changes in speciation/extinction. Finally, we present an overview of both classic and novel hypotheses for why poricidal flowers evolve and suggest directions for future research about the extent to which convergence in poricidal flower morphology is accompanied by convergence in function.

## Material and Methods

### Literature survey

We defined “poricidal flowers” as flowers with tube-like floral structures (typically formed by individual anthers or groups of anthers, but sometimes by staminodes, sepals or corolla), restricting pollen release via an opening that can be described as a pore, small slit, or valve. We conducted our literature search between 2015 and November 2020 for English, German, and Portuguese language articles describing all presently known genera of functionally poricidal plant species. Sources included online books, online search engines (Web of Science, Google Scholar), dissertations, and primary research papers. Search terms included “anther”, “stamen”, or “flower” and “apical pore”, “dehisce”, “dehiscence”, “dehiscent”, “introrse”, “longitudinal”, “porandrous”, “porate”, “pore”, “pored”, “poricidal”, “porose”, “porous”, “short slit”, “thecal pores”, “valve”, or “valvular”. The search was performed at the genus level because numerous well-characterized genera are monomorphic for the presence/absence of poricidal morphology (e.g., *Solanum*, *Senna*, *Miconia*) and comprehensive species-level information for rarer and/or tropical genera is frequently sparse (although we also recognize that monomorphism within genera should not be taken for granted and there are exceptions in some genera, e.g., Renner, 1989; Gavrutenko *et al*., 2020). For each genus, we documented the flower parts involved in forming the tubular structures that comprise poricidal flower morphology, as well as how pollen is released from poricidal anthers (Fig. S2, Table 1; Supplementary Material Table S1). We also confirmed non-poricidal morphology for monomorphic genera when morphological information was available, but generally did not catalog the accompanying literature.

**Table 1.**
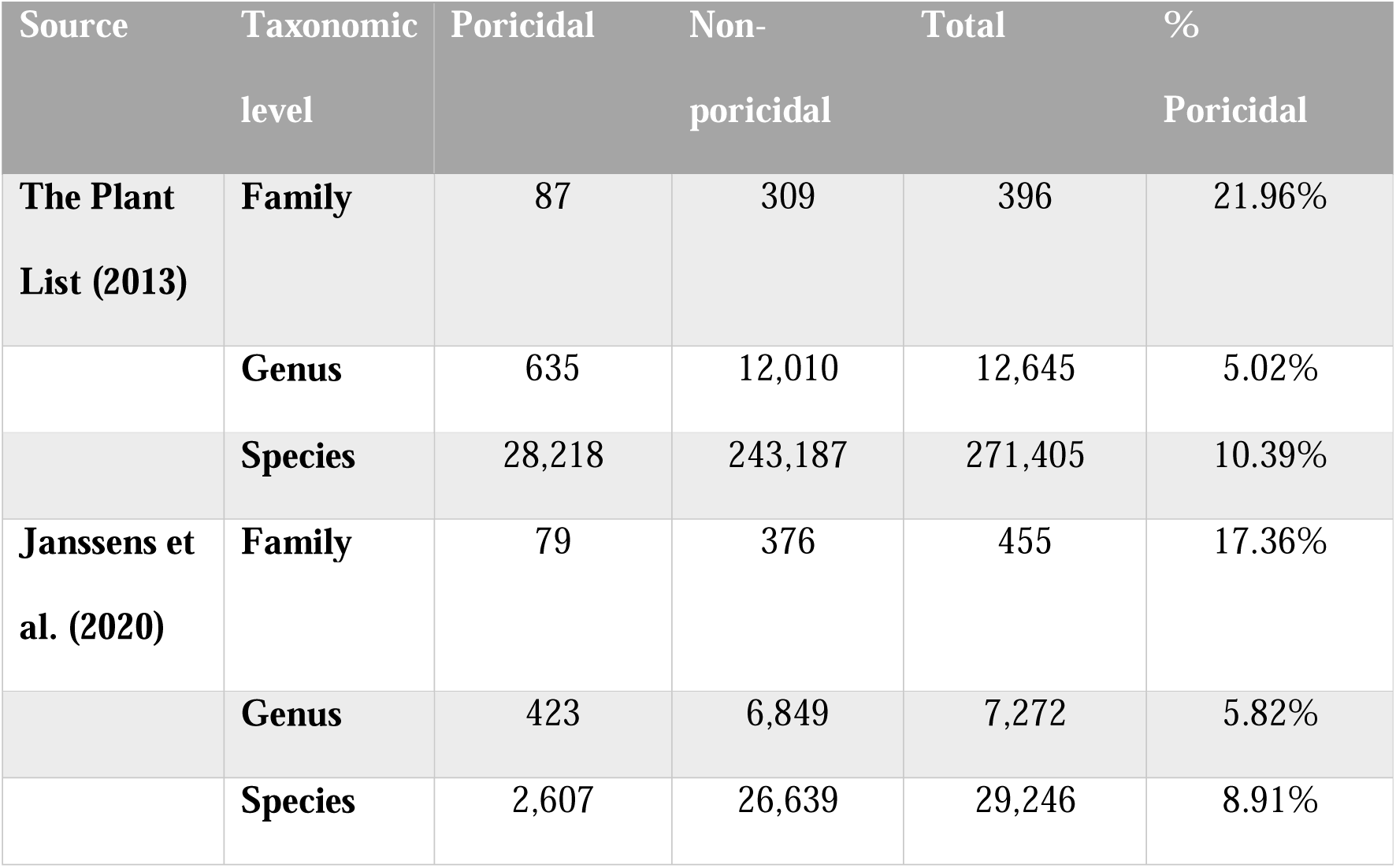
Incidence of poricidal flowers at the species, genus, and family levels, as estimated from an angiosperm-wide database and within the Janssens et al. (2020) phylogeny used for the angiosperm-wide analysis. The proportions of poricidal flowers at each taxonomic level are similar. Poricidal = number of taxa in which at least one species or genus is reported as poricidal (genera treated as monomorphic). Non-poricidal = Number of taxa with no reported poricidal species or genera.

| Source | Taxonomic level | Poricidal | Non-poricidal | Total | % Poricidal |
| --- | --- | --- | --- | --- | --- |
| **The Plant List (2013)** | **Family** | 87 | 309 | 396 | 21.96% |
|  | **Genus** | 635 | 12,010 | 12,645 | 5.02% |
|  | **Species** | 28,218 | 243,187 | 271,405 | 10.39% |
| **Janssens et al. (2020)** | **Family** | 79 | 376 | 455 | 17.36% |
|  | **Genus** | 423 | 6,849 | 7,272 | 5.82% |
|  | **Species** | 2,607 | 26,639 | 29,246 | 8.91% |

### Angiosperm phylogeny and estimates of species number per genus

To estimate the incidence of poricidal and non-poricidal species across the angiosperms we cross-referenced our 639 poricidal genera against The Plant List (2013), an angiosperm-wide database, which contained 271,405 species, 12,645 genera, and 396 families of angiosperms, after filtering out infraspecific classifications (i.e., subspecies and varieties). We then matched the Janssens et al. (2020) phylogenetic tree to our species-level data using the R package *treeplyr* (Uyeda and Harmon 2020) and found direct matches for 28,980 species. The Janssens et al. (2020) phylogenetic tree was specifically developed to address species-level macroevolutionary analyses like those performed in the present study. The Janssens et al. (2020) phylogenetic tree is based on matK (incl. trnK) and rbcL. plastids and has 36,101 taxa representing 8,399 genera and was dated using 56 angiosperm fossils. Additionally, there were 266 species in the Janssens et al. (2020) tree (each representing a different genus) that were absent in our species-level data, but were of genera represented in our species-level data. To maximize information for the phylogeny, we assigned the trait value of each genus in our species-level data to these 266 species in the Janssens et al. (2020) tree, via the R package *phyndr* (Pennell *et al*., 2016). This resulted in an ultrametric and bifurcating phylogenetic tree with a total sample size of 29,246 taxa (tips) with poricidal binary states that we used for the phylogenetic comparative analyses below.

### Phylogenetic comparative methods

#### State dependent diversification and origination analyses

The 29,246 taxa in the resulting time-calibrated bifurcating tree have tips coded as binary states (‘non-poricidal’ or ‘poricidal’) as the input of a hidden state dependent speciation and extinction model (HiSSE, Beaulieu and O’Meara 2016), with the goal of testing whether differences in diversification between poricidal and non-poricidal states exist. For every HiSSE model we identify the differences in net diversification (speciation minus extinction) between poricidal and non-poricidal states, but also the diversification differences given by hidden states A and B. The difference between hidden states is of particular interest, especially in large phylogenies, because it represents the potential for diversification due to unmeasured factors or heterogeneity, but not the trait of interest (Beaulieu & O’Meara, 2016). Thus, estimating the differences between the main and hidden states disentangles the effect of poricidal state on diversification from other potential sources of diversification. We only fitted HiSSE and not a simpler model without hidden states (BiSSE, Maddison *et al*., 2007) for two main reasons. First, in large phylogenetic trees, many unobserved traits likely contribute to diversification, and thus our null hypothesis should always have heterogeneity in diversification rates due to these unobserved traits (instead of the trait of interest having all the signal). Second, in a Bayesian framework the posterior distribution of the HiSSE model can be re-written to represent the null hypothesis determined by the character independent model with two hidden states (CID-2), which describes a model of diversification where the rates change only due to hidden states and not the trait of interest, as described below.

The HiSSE model was fitted under a Bayesian framework within the RevBayes environment (Höhna *et al*., 2016). We obtained the posterior probability densities for every parameter in the model via a Markov chain Monte Carlo (MCMC) algorithm. We checked for convergence of our MCMC algorithm by running two chains and observing that the posterior distributions reached the same range of values, and that stationarity was achieved (see Supplementary Material for details). Convergence via Gelman and Rubin’s convergence diagnostic (Brooks and Gelman 1998) was verified using the software TRACER (Rambaut *et al*., 2018) (Fig. S4-S6). We assumed the sampling fraction for the HiSSE model was 10%, given the current estimate of ∼300,000 extant flowering plant species (The Angiosperm Phylogeny Group 2016). We do not add a sampling correction for states, because our tree reflects our literature survey at the species, genus, and family level (Table 1).

To examine whether there might be family specific patterns in how poricidal state might influence diversification rates, a second round of HiSSE models were performed for four large angiosperm families matched to the Smith and Brown (2018) phylogenetic tree: Solanaceae, Melastomataceae, Ericaceae, and Fabaceae. These families were selected because they have well-resolved trees, are large, and are variable for poricidal state. For each of the four families we pruned the species-level phylogenies from the Smith and Brown (2018) tree and matched them with poricidal status. The matching trees and states on the tips were the inputs for HiSSE. All these analyses were accomplished via RevBayes (Höhna *et al*., 2016) and convergence was checked similarly to our angiosperm-wide phylogenetic tree. We reached an ESS of more than 200 for all parameters.

#### Testing state-dependent diversification in a Bayesian framework

The character independent models (Beaulieu and O’Meara, 2016) are at the core of testing if the trait of interest is linked to diversification. In the approach used by the R package *hisse* (Beaulieu *et al*., 2023) multiple models of state-dependent diversification are fitted to data and then a model selection procedure using the Akaike Information Criterion (AIC) determines if a model that includes diversification rates linked to the main states is preferable over a model with diversification rate differences given only by the hidden states. When fitting state-dependent diversification models in a Bayesian framework, as done in this study, the model selection part of the process is unnecessary since the samples of the posterior distributions for the parameters of HiSSE can be transformed to test the hypotheses laid out by a CID-2 model as we describe here. The null hypothesis in CID-2 assumes that the two main states (non-poricidal and poricidal) are equal within each hidden state (A or B), but a given main state is different between hidden states. In statistical terms, the null hypothesis is:

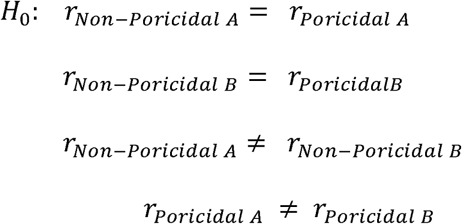

Where *r*_i_ refers to the net diversification (the difference between speciation and extinction, in mathematical terms, A_i_ - µ_i_) in state i. Using our species-level HiSSE results (Fig. 2b *H*_0_), we created two test statistics to determine the probability of the differences between non-poricidal and poricidal states within each hidden state. These test statistics represent the differences of net diversification between non-poricidal and poricidal states within each hidden state. Mathematically, these can be written as:

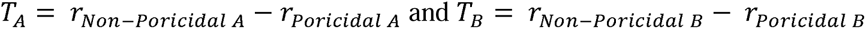

and their posterior distributions are calculated by taking the differences in the speciation and extinction sampling estimates in each iteration of MCMC (Gelman *et al*., 1995). For hypothesis testing, we calculate the quantile that the value of 0 represents (*T*_A_ =0): the probability of the absolute value of T_A_ being greater than zero (in mathematical notation *P*(|*T_A_*|> 0)). If *P*(|*T_A_*|> 0) > 0.05, non-poricidal and poricidal states in hidden states A are the same, similarly for *T_B_* =0. This simple test statistic transformation negates the need for model selection, while allowing us to test for the same hypothesis delineated by CID-2.

**Figure 2.**
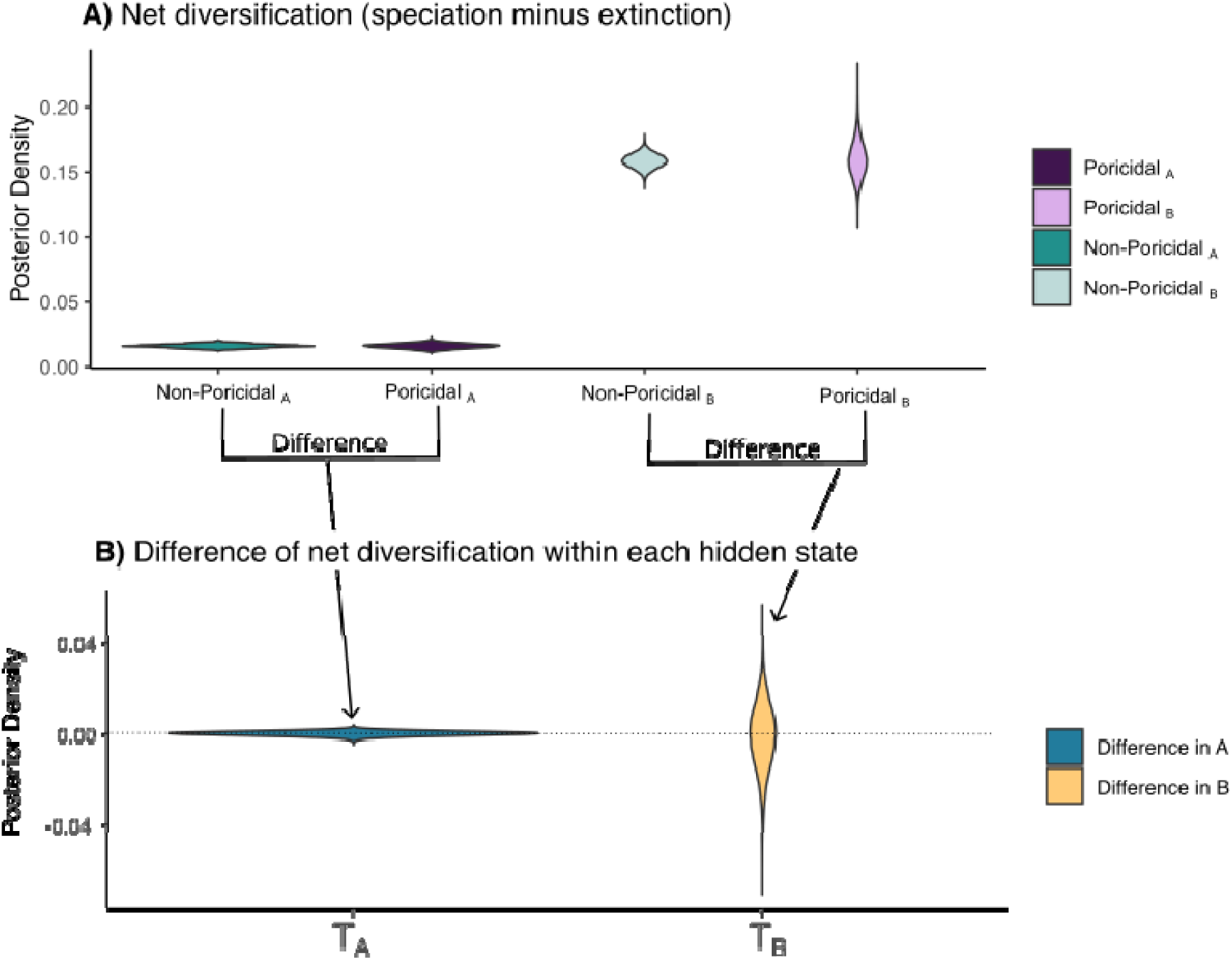
A) Posterior distribution of net diversifications under the hidden state dependent speciation and extinction model (HiSSE) for non-poricidal and poricidal states. No differences in diversification are found between main states, but only in hidden states. B) Test statistics and that calculate the difference between poricidal and non-poricidal states in each hidden state show that it is highly probable that there are no differences in diversification due to poricidality.

**Figure 3.**
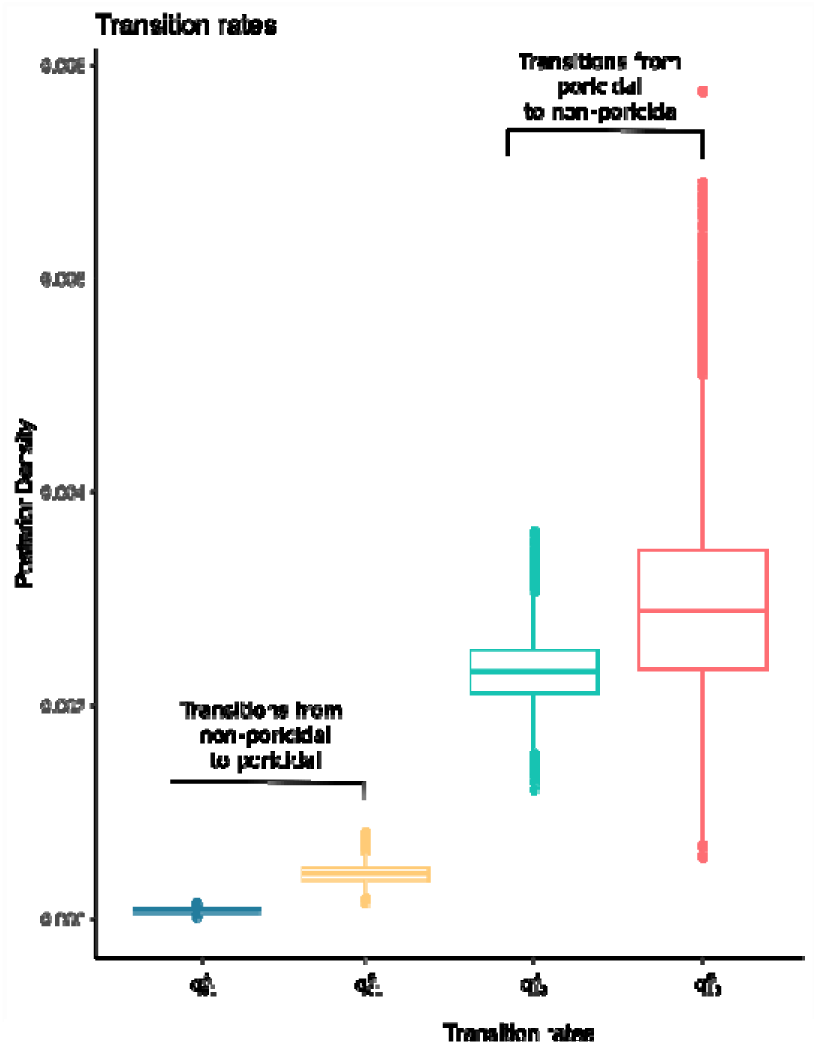
Posterior distribution of rates of transition between non-poricidal and poricidal states under hidden states A and B using the hidden state dependent speciation and extinction model (HiSSE). For both hidden states it is easier to transition out of the poricidal state than into the poricidal state ( ).

#### Models without diversification. Estimation of number of poricidal transitions and their timing

As discussed in the Results section, we found that poricidal status does not inform the process of diversification. For this reason, we fitted a simpler model: a Markov model with two states (poricidal or not poricidal) and hidden states (A and B), excluding diversification rates. We estimated the transition rates of this model using the R package *corhmm* (Beaulieu *et al*., 2022). Under this model, we estimated 1000 stochastic maps using R package *phytools* (Revell, 2012) to extract the distribution of the expected number of transitions between poricidal and non-poricidal states in both hidden categories, and to characterize the distribution of waiting times where the transitions between states happened. The distribution of the number of transitions allows us to assess which transitions are more common, and the distribution of waiting times shows whether these transitions are more common at specific times in the evolution of angiosperms. We calculate the relative lineage through time plots per state by calculating the lineages through time possessing a given state, and at a given time interval, by dividing them by the total lineages. This relative LTT allows us to compare differences in the distributions of lineages at a given time point that are difficult to observe in a 29K-tip phylogenetic tree (e.g., two lineages at 199 mya vs. 29K lineages at 0 mya) (Fig. 5).

## Results

### Distribution of poricidal flowers across taxonomic levels

In a seminal paper, Buchmann (1983) estimated that 15,000-20,000 species of angiosperms in 72 families and 544 genera have anthers that dehisce through apical pores or slits (poricidal anthers). Here, we found that poricidal flowers, including species with poricidal anthers, are reported from 87 families and 639 genera (Fig. S2, Table 1; Table S1). Using the 271,405 species of angiosperms in the The Plant List database, which includes 635 of 639 genera identified in our study, yields an upper estimate of 28,218 poricidal species (21.96% of plant families, 5.02% of genera, and 10.39% of species) (Table 1). Our results therefore suggest that poricidal flower morphologies have evolved in a fifth of families and up to one in ten angiosperm species.

### Different types of poricidal flowers across taxonomic levels

Flower parts forming the tubular structures that comprise poricidal flower morphology include poricidal anthers (an estimated 97% of poricidal species), corollas, or other functionally poricidal floral parts (i.e., staminodes or sepals) (Table 2). Additionally, how pollen is released from poricidal anthers is particularly diverse, although such classifications are morphological, rather than developmental, and may therefore be quite inaccurate. Morphological variation includes pores (60 families and 441 genera), short terminal slits (46 families and 125 genera), valves (15 families and 74 genera), and introrse dehiscence forming terminal pores or short slits (16 families and 53 genera) (Supplementary Material Table S1). A given poricidal flower morphology is rarely restricted to a particular family or genus and even multiple kinds of poricidal anthers are known within a genus (Endress, 1996).

**Table 2.**
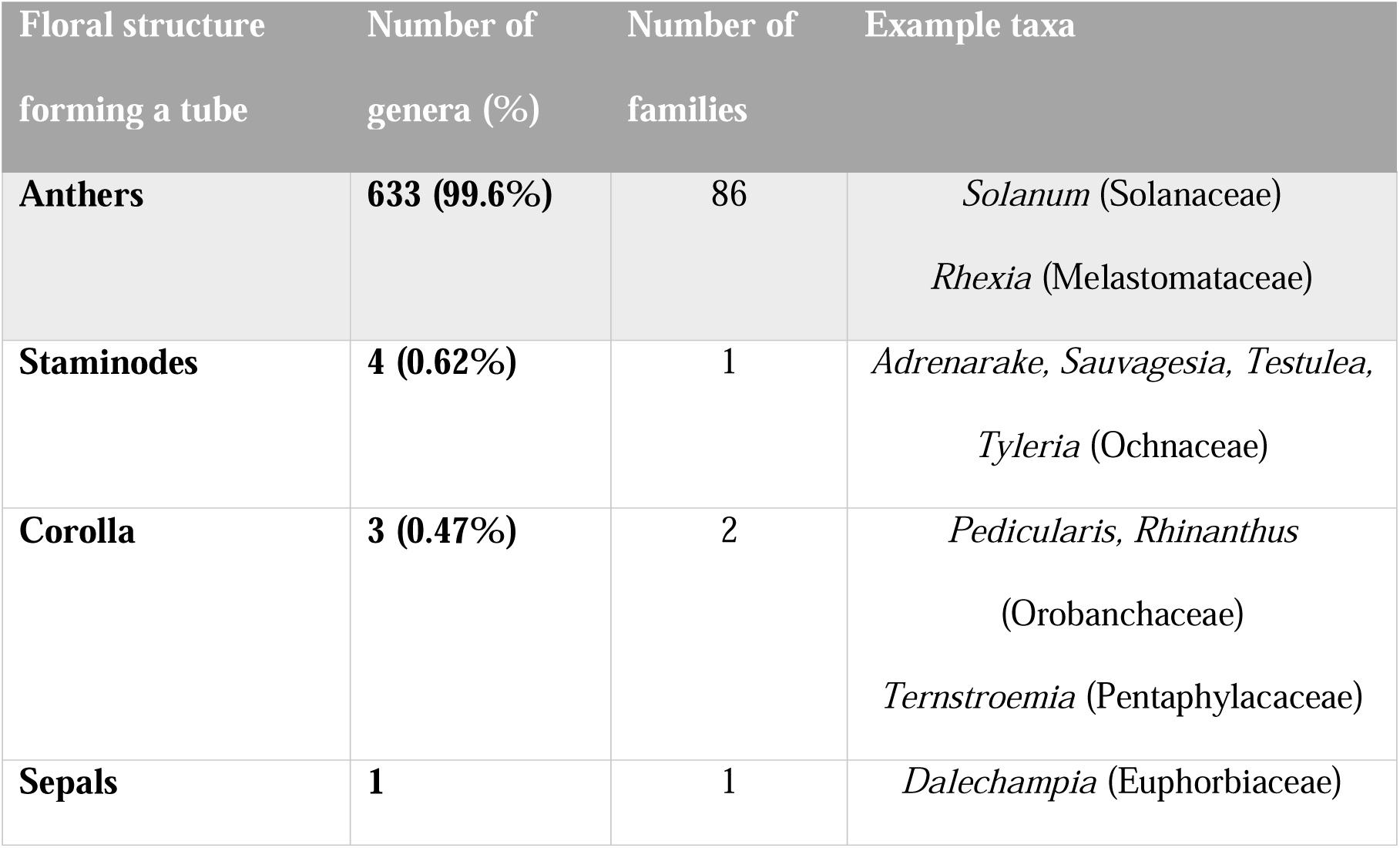
Type of floral structure involved in forming a tube-like structure with a small opening containing pollen grains (poricidal flower) exhibited by 639 genera. Two genera (*Tyleria* and *Testulea*; Ochnaceae) contain species forming poricidal flowers via pores and staminodes and thus are included more than once in the table. Data from Supplementary Material Table S1.

### Diversification rates and transitions between poricidal and non-poricidal flowers

For the angiosperm-wide species-level HiSSE analyses using the (Janssens et al. 2020) phylogenetic tree with our data, we found that there were no differences in net diversification (speciation minus extinction rates) between non-poricidal and poricidal states (Fig. 2). Instead, we only found differences between hidden state net diversification rates with probability (*P*(|*T_A_*|> 0) =0 and *P*(|*T_B_*|> 0) = 0) (Fig. 2B). This result is consistent with the synthesis on state-dependent diversification presented in Helmstetter et al. (2023), where discrete traits modeled under SSE approaches in trees with large numbers of tips typically have no correlation with diversification rates. In the species-level Solanaceae HiSSE analysis, poricidal flowers diversified at a higher rate compared to non-poricidal flowers, but again, this relationship varied with the state of the hidden character. In the HiSSE analysis of Ericaceae, we found a higher diversification rate of poricidal flowers in combination with hidden character state A, but lower when combined with character state B, indicating a strong context-dependence effect of poricidal morphologies (Fig. S2). We lack results for the two other families analyzed at the species level (Melastomataceae and Fabaceae), because models did not converge after running the analyses for several months, due to high uncertainty on the branch lengths (additional discussion in Supplementary Material).

Our diversification results for the angiosperm-wide species-level HiSSE analysis indicate that the evolution of poricidality is faster for certain lineages (in hidden state B), but not for all angiosperms. Furthermore, we found that transitions from non-poricidal to poricidal states are on average 33x and 6.6x slower than from poricidal back to non-poricidal in A and B hidden states, respectively (95% credible intervals: non-poricidal to poricidal: ), and ); poricidal back to non-poricidal: ), and )), (Fig. 5B). This pattern of asymmetry in transition rates from a “specialized” poricidal state back to a “generalized” non-poricidal state was also found in recent state-dependent diversification analyses (Day *et al*., 2016; Zenil-Ferguson *et al*., 2023), where more evidence of evolving out of ecological specialization has been found.

### Discrete trait evolution recovers asymmetry of transitions outside of poricidal state

After fitting a discrete state model with hidden states, we also recover asymmetry in maximum likelihood estimates (q_01_^A^ = 4.68x10^-3^, q_01_^B^ = 2.34x10^-6^, q_10_^A^ = 1.28x10^-2^, and q^B^ = 2.19x10^-4^). The software would not permit recovering likelihood or confidence intervals for these maximum likelihood estimates and thus we cannot test if the estimates are significantly different. However, it is worth noting that point estimates differ by at least 100x. Using the 1000 stochastic maps simulated from the discrete and hidden states Markov model (Mkn), we calculated the distribution of the expected number of changes between states. In Table 3 we summarize the statistics of these transitions, and found the same pattern, with fewer transitions into poricidal states than transitions out of poricidal states.

**Table 3.** Summary statistics of transitions obtained from the 1000 stochastic maps. These statistics were obtained by following each lineage and counting the number of transitions from one state to the next.

| Type of transition | Mean | Median | Credible Intervals (95%) |
| --- | --- | --- | --- |
| Non-Poricidal<sub>A</sub> →<br>Poricidal<sub>A</sub> | 202.22 | 202 | (184, 220) |
| Poricidal<sub>A</sub> →<br>Non-Poricidal<sub>A</sub> | 214.79 | 215 | (189, 239) |
| Non-Poricidal<sub>B</sub> →<br>Poricidal<sub>B</sub> | 3.75 | 4 | (1, 7) |
| Poricidal<sub>B</sub> →<br>Non-Poricidal<sub>B</sub> | 22.85 | 23 | (15, 31) |

### Timing of poricidality: A fifty million year wait and establishment

Under the discrete and hidden states Markov model (Mkn) we simulated 1000 stochastic maps, but we present just one (Fig. 4). In this figure, we observed that poricidal states (Poricidal_A_ (red) and Poricidal_B_ (yellow)) evolved around 150 million years ago (∼ 50 million years after the origin of angiosperms; Fig. 5) and remain in that state until the present (especially in state Poricidal_B_, Fig. 4). This observation is reinforced by the relative lineage through time plots (Fig. 5c and 5f), where state Poricidal_A_ suddenly increases in relative numbers, whereas Non-Poricidal_A_ decreases. Later, state Non-Poricidal_A_ mostly transitions into Non-Poricidal_B_, and Poricidal_A_ into Poricidal_B_ and grows in numbers without switching further states, as seen in the single stochastic map (Fig. 4).

**Figure 4.**
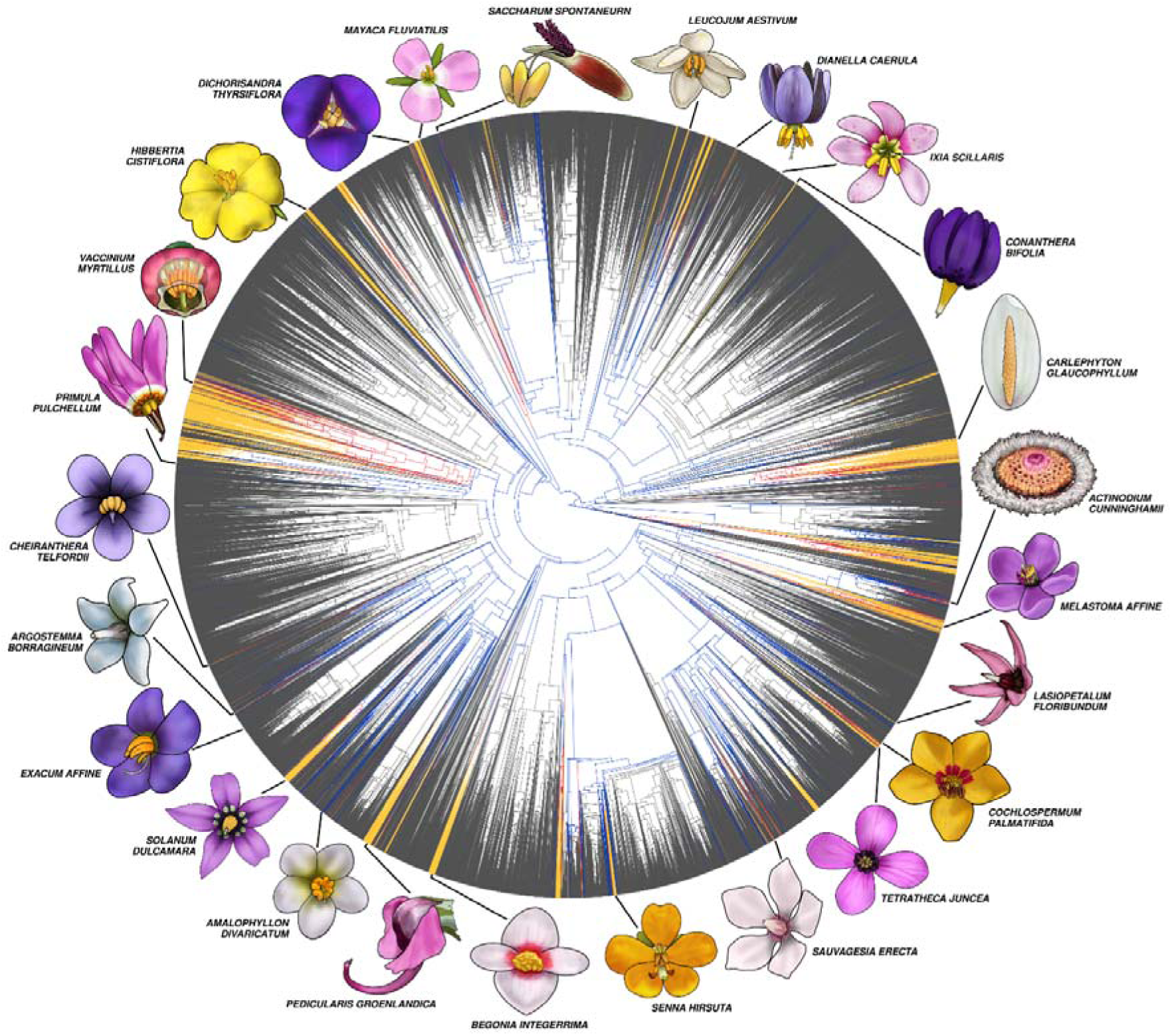
A single representative stochastic map calculated via an angiosperm-wide phylogenetic tree with 29,246 taxa under a discrete-state and hidden state model using R packages *corhmm* (Beaulieu et al. 2022) and *phytools* for visualization (Revell 2012). Non-poricidal states in blue (0A) and grey (0B). Poricidal states in red (1A) and yellow (1B). Even when non-poricidal states (light gray) represent the majority of the diversification in angiosperms, some yellow “bursts” of poricidal state happened, such as within large families like Solanaceae, Ericaceae, Fabaceae, and Melastomataceae. Digital art of illustrative poricidal flowers by Moth Castagna.

**Figure 5.**
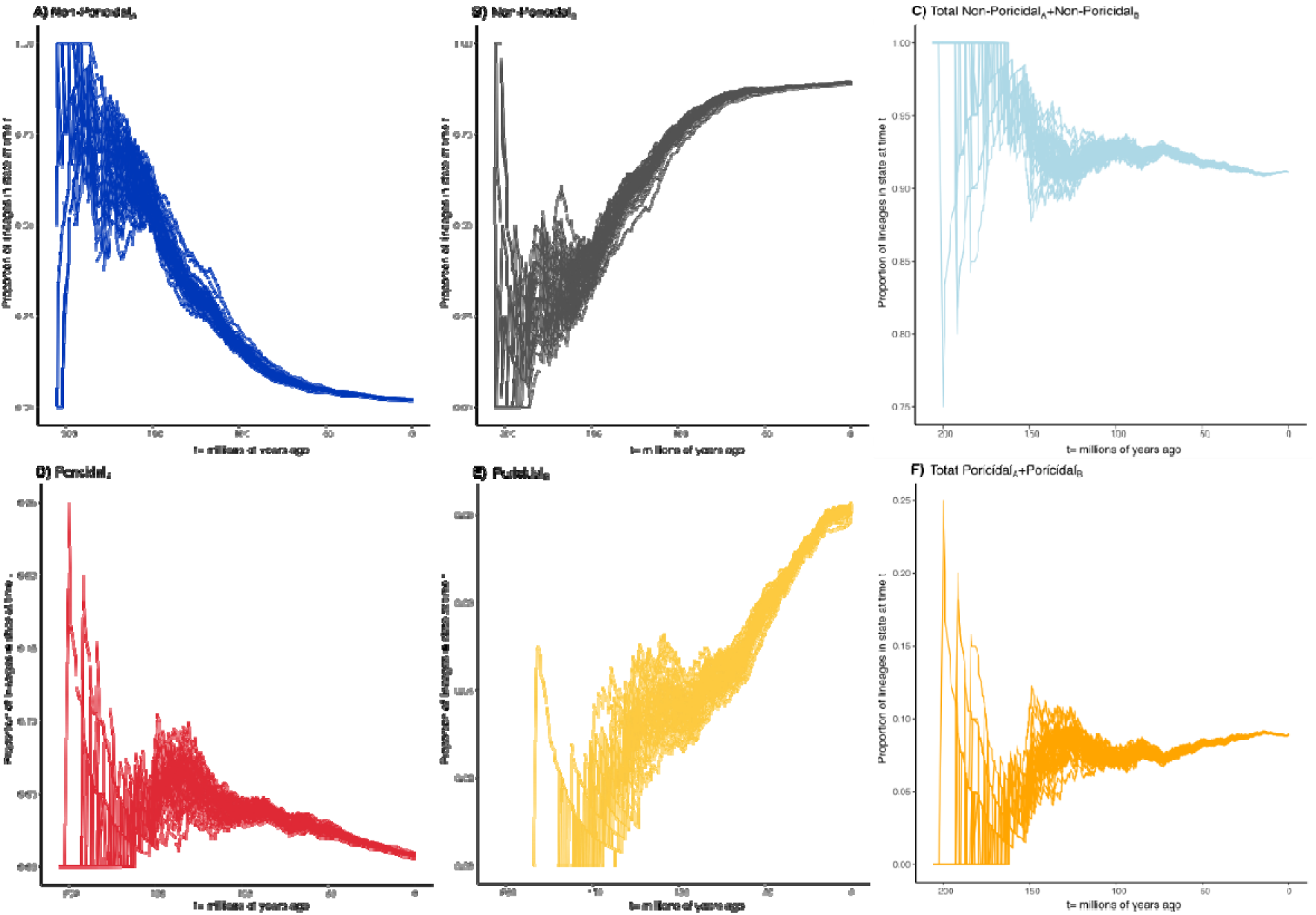
Relative lineage through time (RLTT) plots from the 1000 stochastic maps. The relative lineage through time plots are calculated by counting the number of lineages through time in each state and dividing them by the total number of lineages. A) The RLTT for Non-Poricidal_A_ state indicates a decline of lineages in this state starting around 180 million years ago, with this state mostly transitioning into B) Non-Poricidal_B_ state and D) Poricidal_A_. F) Poricidal_A_ state transitions later into Poricidal_B_. C) and F) have the total in each main state shown independently from the hidden state.

## Discussion

Our study shows that the repeated evolution of functionally poricidal floral morphology represents a major case of convergent evolution in flower form across angiosperms. Although we found no overall effect on speciation-extinction rates, effects of evolving poricidal flowers on species diversification likely strongly depend on the phylogenetic, and probably ecological, context in which they occur, as suggested by our focal family-level analyses. Our survey updates previous estimates of the incidence of taxa with functionally poricidal floral morphology, showing that about 10% of angiosperm species have this morphology. We can also provide a quantitative estimate of its repeated evolution, with an average minimum of 205 independent origins across the angiosperms (Table 3). Although poricidal flowers have repeatedly independently evolved, which selective pressures drive their evolution are currently unknown. Given the diversity of angiosperm clades and ecosystems in which poricidal flowers have arisen (Buchmann 1983; Russell *et al*., 2024), and the multiple non-mutually exclusive hypotheses explaining their overall evolution (Vallejo-Marin & Russell, 2024), comparative studies across multiple independent origins of poricidal flowers are needed to identify the role of different selective pressures on the evolution of this convergent floral morphology.

When considering angiosperms as a whole, we find significantly lower origination rates for poricidal taxa (33x - 6.6x lower), suggesting that this trait is frequently costly. One obvious potential cost is that poricidal morphology increases pollination specialization, reducing the capacity for these plant taxa to persist or diversify (Renner, 1989; Dellinger *et al*., 2019b). Specialization in a variety of ecological systems (e.g., parasitism, herbivory, pollination) can be associated with reduced diversification rates (Moran, 1988; Tripp & Manos, 2008; Day *et al*., 2016; Raia *et al*., 2016) although this relationship may not be straightforward (Forister *et al*., 2012; Armbruster, 2014; Hardy & Otto, 2014; Day *et al*., 2016; Zenil-Ferguson *et al*., 2023). At the same time, specialization is easy to reverse to a generalist non-poricidal state (Fig. 3) and poricidal flowers might be costly to reacquire, which could explain why poricidal states are not linked to recent diversifications for most angiosperms (Fig. 5D). A main reason for the difficulty in reacquiring the poricidal state is that animal pollination in poricidal plant species is frequently restricted to a subset of possible pollinators. In fact, many poricidal plant taxa are pollinated by bees capable of floral buzzing or, rarely, by birds or bees that activate a bellows-like mechanism in the stamen (Sazima *et al*., 1993; Dellinger *et al*., 2019b) and is likely often more specialized. Furthermore, the regional diversity of poricidal taxa and buzzing bees can be strongly positively correlated (Pacheco Filho *et al*., 2015), suggesting an even greater degree of poricidal plant-bee specialization (MesquitaLNeto *et al*., 2018).

The costs of poricidal morphology may also be related to pollen being the only reward offered by the majority of poricidal species (Vogel, 1978; Buchmann, 1983; Endress, 1994; Vallejo-Marin *et al*., 2010). In addition to resulting in specialization on pollen-foraging pollinators, pollen offered as a food reward might reduce its availability for export to conspecific flowers (Hargreaves *et al*., 2009). Furthermore, while separating male and female phases in flowers (dichogamy) reduces selfing and/or interference with cross-pollinating mechanisms, dichogamy in pollen-only rewarding species also potentially reduces visitation to the rewardless female phase flowers. If the costs of dichogamy outweigh its benefits for pollen-only rewarding species, this may explain why the simultaneous presentation of pollen and stigmas (homogamy) in pollen-only flowers is common (Lloyd & Webb, 1986; Webb & Lloyd, 1986; Renner, 1989).

Notwithstanding the various potential costs of poricidal morphology and its associated traits, some plant families have higher diversification rates associated with poricidal flowers, although this is contingent on the state of hidden characters (e.g., Solanaceae, Ericaceae). These hidden characters might for instance include the availability of the right species of buzz-pollinating bees or optimal environmental conditions for facilitating pollen while being vibrated (e.g., low aridity and low wind; see Russell *et al*., 2024; Vallejo-Marin & Russell, 2024). Other factors that have been suggested to modulate the diversification of poricidal taxa include flower modularity (Dellinger *et al*., 2019a), the relative availability of potential pollinators (Pacheco Filho *et al*., 2015; MesquitaLNeto *et al*., 2018), or the presence of other pollen sources in the community (Kemp *et al*., 2022).

Poricidal morphology is thought to be an adaptive plateau and appears to constrain floral evolution (Renner, 1989; Dellinger *et al*., 2019b). Yet poricidal floral morphology is not necessarily a dead end. We estimate 215 independent losses of this morphology on average, suggesting this trait is evolutionarily labile. Multiple non-mutually exclusive hypotheses might explain a return to loculicidal anther dehiscence. For example, the loss of key pollinators, a reduction in herbivory, and/or reduced selection by microbial pathogens could all reduce the potential benefits of poricidal versus non-poricidal morphology (Schwartz-Tzachor *et al*., 2006; Brito *et al*., 2017). A detailed understanding of the biogeography of poricidal taxa and of pollinators and other selective agents will likely be required to distinguish among these hypotheses (see Russell *et al*., 2024), which the present dataset will facilitate. Given differences in how functional poricidal morphology is achieved, certain poricidal types may also be more evolutionarily labile than others. For example, the length of the distal slits in poricidal anthers often varies among taxa or even over the lifetime of a given flower (e.g., in *Actinidia*; Goodwin, 1986), and thus evolutionary transitions from partial to more and more complete longitudinal dehiscence for some floral forms may be developmentally relatively simple to achieve.

Finally, we catalogue substantial diversity and bias in how poricidal morphology is achieved (Fig. 1; Table 2). While floral parts such as anthers, corollas, sepals, and/or staminodes can be involved, anther modifications are particularly common (∼99% of poricidal genera Table 2, S1). Within taxa, variation in which floral parts are used to achieve poricidal morphology is uncommon. Additionally, there is substantial morphological diversity in poricidal anther among taxa, especially with respect to anatomical differences in dehiscence (e.g., via slits, pores, or valves; Harris, 1905; Buchmann & Hurley, 1978; Buchmann, 1983; Endress, 1996). These patterns strongly suggest developmental biases, but the developmental pathways of poricidal morphology remain unexplored to our knowledge. Our study sets the stage for future work to investigate the evolutionary history of floral traits that repeatedly proposed to be potentially correlated with poricidal floral morphology, such as loss of nectar, pore size, pollen size and exine ornamentation, pollen quantity, or dry and loose pollen (Vogel, 1978; Dukas & Dafni, 1990; Cruden, 2000; Roulston *et al*., 2000; Freitas & Sazima, 2003; Brito *et al*., 2016).

Classical hypotheses for the evolution of poricidal morphology have highlighted the role of pollinators (Vallejo-Marin & Russell, 2024). Most, but not all species with poricidal flowers are pollinated by floral buzzing bees, which are prolific consumers of pollen (Michener, 1962; Buchmann, 1983; Hargreaves *et al*., 2009; Danforth et al. 2019; Vallejo-Marin, 2022). Poricidal morphology might control the rate of pollen release, and thus reduce the tempo of pollen removal by pollen-consuming pollinators (e.g., Larson & Barrett, 1999; De Luca *et al*., 2014; Brito *et al*., 2016). Likewise, poricidal morphology may result in more effective dispersal of pollen to conspecifics by pollinators, by enhancing the precision of pollen placement on the pollinator body and/or by filtering more generalized and less effective pollinators (e.g., Pacheco Filho *et al*., 2015; MesquitaLNeto *et al*., 2018; Kemp & Vallejo-Marin, 2021; Vallejo-Marin & Russell, 2024). However, pollinator-mediated selection alone appears insufficient to explain the evolution of poricidal morphology across diverse angiosperm clades. Poricidal species have diverse mating systems, from selfing to outcrossing, and many poricidal flowers are wind pollinated, not animal pollinated (e.g., in Poaceae and Halophytaceae; Pozner & Cocucci, 2006; Table S1). Indeed, some poricidal taxa arose as early as 150 million years ago, potentially ∼26 million years before buzz pollinating bees (Cardinal *et al*., 2018; Almeida *et al*., 2023). Although vibration can be a key mechanism of wind-mediated pollen release (Timerman & Barrett, 2018), and could function analogously to floral buzzing by bees for these poricidal species, the functional benefits of poricidal morphology likely go well beyond pollen release, as discussed in Vallejo-Marin and Russell (2024).

In conclusion, given the hundreds of thousands of non-poricidal taxa and the tens of thousands of poricidal taxa, transitions to poricidal morphology are relatively rare, but once evolved, the probability of losing poricidal morphology is relatively high. Other possibilities for this pattern include early bursts of speciation associated with poricidal morphology or greater chances for the poricidal state to go extinct later, given that poricidal genera are proportionally more common early in angiosperm evolution, but later become rarer relative to non-poricidal genera. Our results thus suggest that selective forces driving the evolution and maintenance of poricidal morphology may vary substantially over time and space. Given our coding of genera as poricidal or not, our estimates for the independent loss and gain of poricidal morphology are also likely highly conservative and should be interpreted cautiously. Despite extensive documentation of poricidal morphology (Supplementary Material Table S1), detailed data across multiple species has only been collected for some of the major lineages (Brito *et al*., 2016; Dellinger *et al*., 2019a; Dellinger *et al*., 2019b).Furthermore, the pollinators or even pollination systems of poricidal species are frequently incompletely documented. Additionally, poricidal flowers are associated with diverse pollination systems, including many wind-pollinated Poaceae, fly and/or non-buzzing bee pollinated Araceae, Rafflesiaceae (taxa with toothpaste-like pollen extrusion), and *Rhododendron* (taxa with clumped pollen attached via sticky viscin threads), and bird pollinated *Agarista* and *Axinaea* (taxa with bellows flowers). Considering such diversity, future studies investigating functions beyond only the control of how and when pollen is released will be invaluable in understanding the selective pressures driving the gain and loss of poricidal morphology.

## Data Accessibility

The datasets and code supporting this article will be uploaded to Dryad and made publicly available upon publication.

## Acknowledgements

This paper is dedicated to the memories of James A. Harris, Charles D. Michener, and Stefan Vogel for their foundational efforts in elucidating the ecology of poricidal flowers and their bee pollinators.

