## Supplementary Material for "Widespread evolution of poricidal flowers: A striking example of morphological convergence across flowering plants"

**Running title**: Convergent evolution of poricidal flowers

### **Hidden state-dependent speciation and extinction for phylogenies for families**

We ran hidden state-dependent speciation and extinction models (HiSSE) for four large families of angiosperms: Solanaceae, Ericaceae, Melastomataceae, and Fabaceae.

The results for Solanaceae and Ericaceae are shown in Fig. S1.

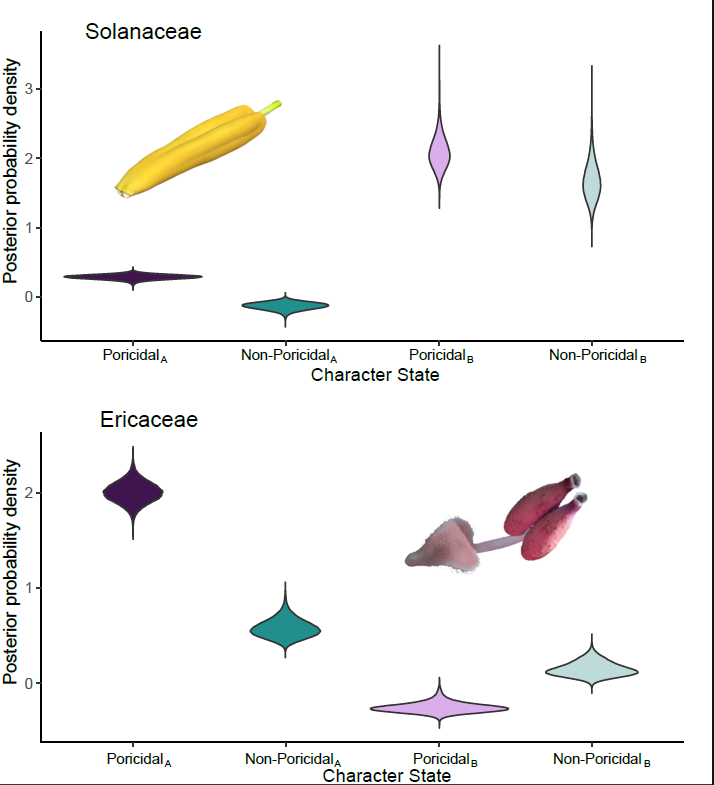

***Figure S1.*** *Posterior probability density of estimated diversification rates (speciation – extinction) of poricidal vs. non-poricidal taxa (species) in HiSSE models with two hidden states (A, B) in two families: Top panel = Solanaceae; Bottom panel = Ericaceae.*

For the family analyses we chose four families from the Smith and Brown (2018) phylogenetic tree (Solanaceae, Melastomataceae, Ericaceae, and Fabaceae) that are known for high incidence of poricidal species. We pruned them according to the species that had tip information and fitted a HiSSE for each of the families. We did not use any more clade-specific phylogenies, but we crosschecked with the Solanaceae phylogeny of Särkinen *et al.* (2013) and a recent Fabaceae phylogeny of Delaney & Igić (2022), and both of them were analogous in data and ages of the trees; however, no comparisons of family-level phylogenies are available for Ericaceae and Melastomataceae.

Finding accurate species-level phylogenies for other families of angiosperms was extremely challenging. Most phylogenies built to date focus on specific subgroups (genera) of families, due to the sheer size of molecular information that would be needed to create those phylogenies. In addition, most comparative analyses addressing diversification in multiple clades (for example, the ploidy states study from Mayrose *et al.*, 2011) do so for multiple genera, but with two important caveats. First, these analyses create quick and simple phylogenies that aren’t properly time calibrated (for example, Drori *et al.*, 2018). Second, these analyses ignore recommendations for sample size for state dependent-diversification models (Davis *et al.*, 2013 recommended 300 tips, and Mynard *et al.*, 2023 recommends sampling above 50% of taxa for each phylogeny). We tried and failed multiple times at convergence of the models for Fabaceae and Melastomataceae. We think this failure might be due to issues with branch lengths below genus level from super trees, hence the results for only two families.

**Testing for diversification linked to states in a Bayesian framework**

Traditionally, testing for diversification linked to states requires a formal model selection approach as discussed in Beaulieu and O’Meara (2016). The reason for using model selection is because models originally were fitted using a likelihood approach, and in this case an information criterion is necessary to compare between two models evaluated in the maximum likelihood estimates for parameters. When fitting state-dependent diversification models in a Bayesian framework, as done in this study, this process is not necessary because fitting a HiSSE allows us to test directly for the hypothesis that the character independent model (CID) model requires. The following example shows how the approach we used is appropriate to test the hypotheses that character independent models (in particular the CID-2) represent:

The null hypothesis that CID-2 represents argues that the two main states (non-poricidal and poricidal are equal) and that all diversification comes from hidden states (A and B are different). In statistical terms the null hypothesis is:

$$H_{0}: r_{Non-Poricidal A}= r_{Poricidal A}$$

$$r_{Non-Poricidal B}= r_{PoricidalB}$$

$$r_{Non-Poricidal A}\neq r_{Non-Poricidal B}$$

$$r_{Poricidal A} \neq r_{Poricidal B}$$

Using our results of HiSSE for species level (Figure 6 main manuscript) we can quickly test $H_{0}$ in two steps:

Step 1. Testing differences in the main states:

$r_{Non-Poricidal A}=r_{Poricidal A}$ and $r_{Non-Poricidal B}= r_{Poricidal B}$

We create the following test statistics:

$T_{A}= r_{Non-Poricidal A}-r_{Poricidal A}$ and $T_{B}= r_{Non-Poricidal B}- r_{Poricidal B}$

These statistics represent the differences between non-poricidal and poricidal states. A high probability that the test statistics are zero means that the non-poricidal and poricidal states are the same. In Figure S2 the value zero is centered in the posterior distribution of the $T_{A}$ and $T_{B}$ statistics, meaning that the differences in non-poricidal states and poricidal states have a high probability of being equal. The “equivalent” of a p-value would be the posterior probability quantile ${P(T}_{A}>0)=0.49$.

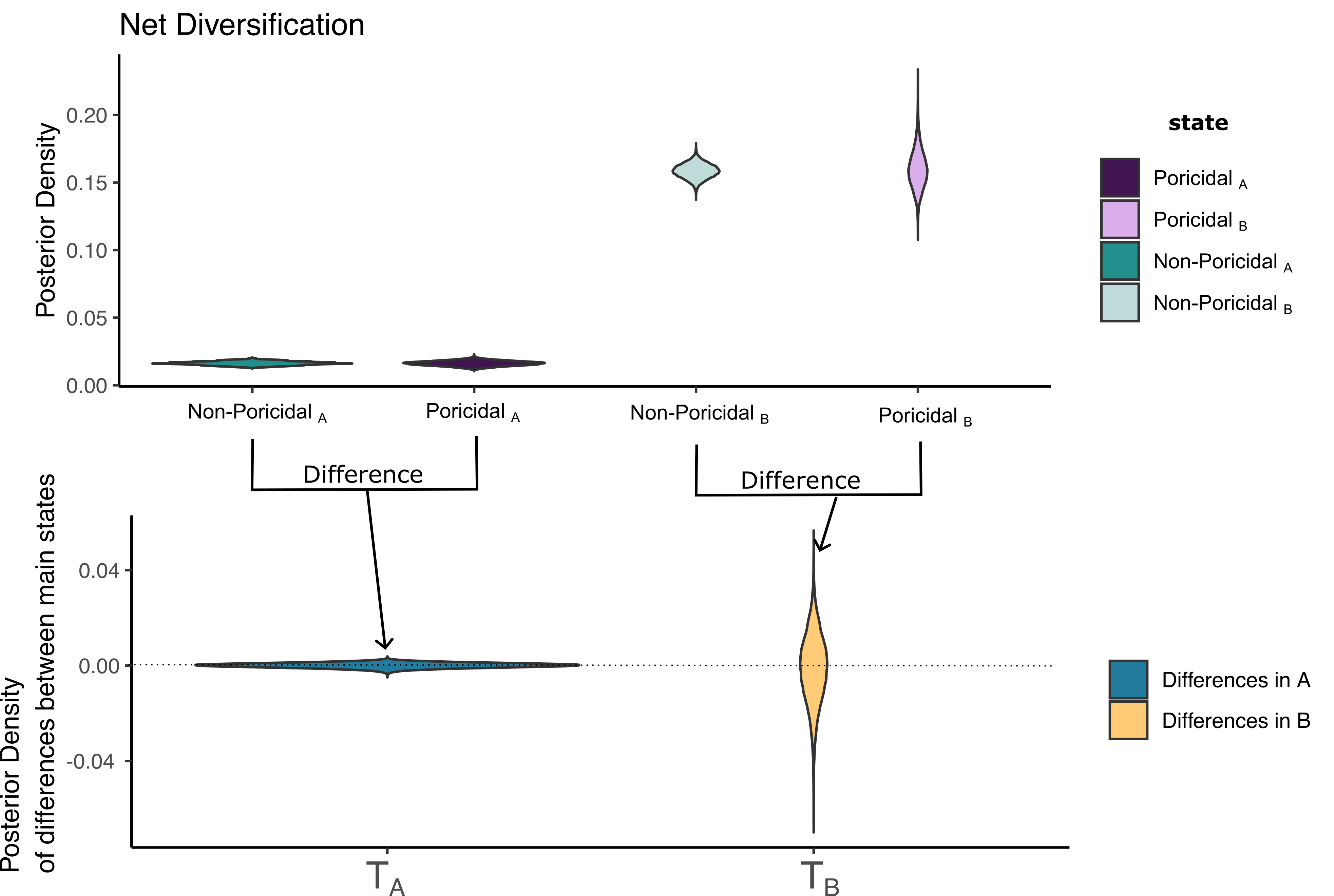

**Figure S2**. Calculating the posterior probability of the test statistics measuring the differences between poricidal and non-poricidal states. The probability for the test statistics$T_{A}$and $T_{B}$ to be zero is high, since zero is located at the center of the probability distribution.

Step 2. Testing the differences between the hidden states:

$r_{Non-Poricidal A}=r_{Non- Poricidal B}$ and $r_{Poricidal A}= r_{Poricidal B}$

We create the following test statistics:

$T_{Non-Poricidal}= r_{Non-Poricidal A}-r_{Non-Poricidal B}$ and $T_{Poricidal}= r_{Poricidal A}- r_{Poricidal B}$

These statistics represent the differences between hidden states A and B. A high probability that the test statistics are zero means that the hidden states are the same. In Figure S3, the value zero is in the right tail in the posterior distribution of these statistics, meaning that the hidden states are different in terms of diversification. The “equivalent” of a p-value would be the posterior probability quantile ${P(T}_{Non-Poricidal}>0)=0$.

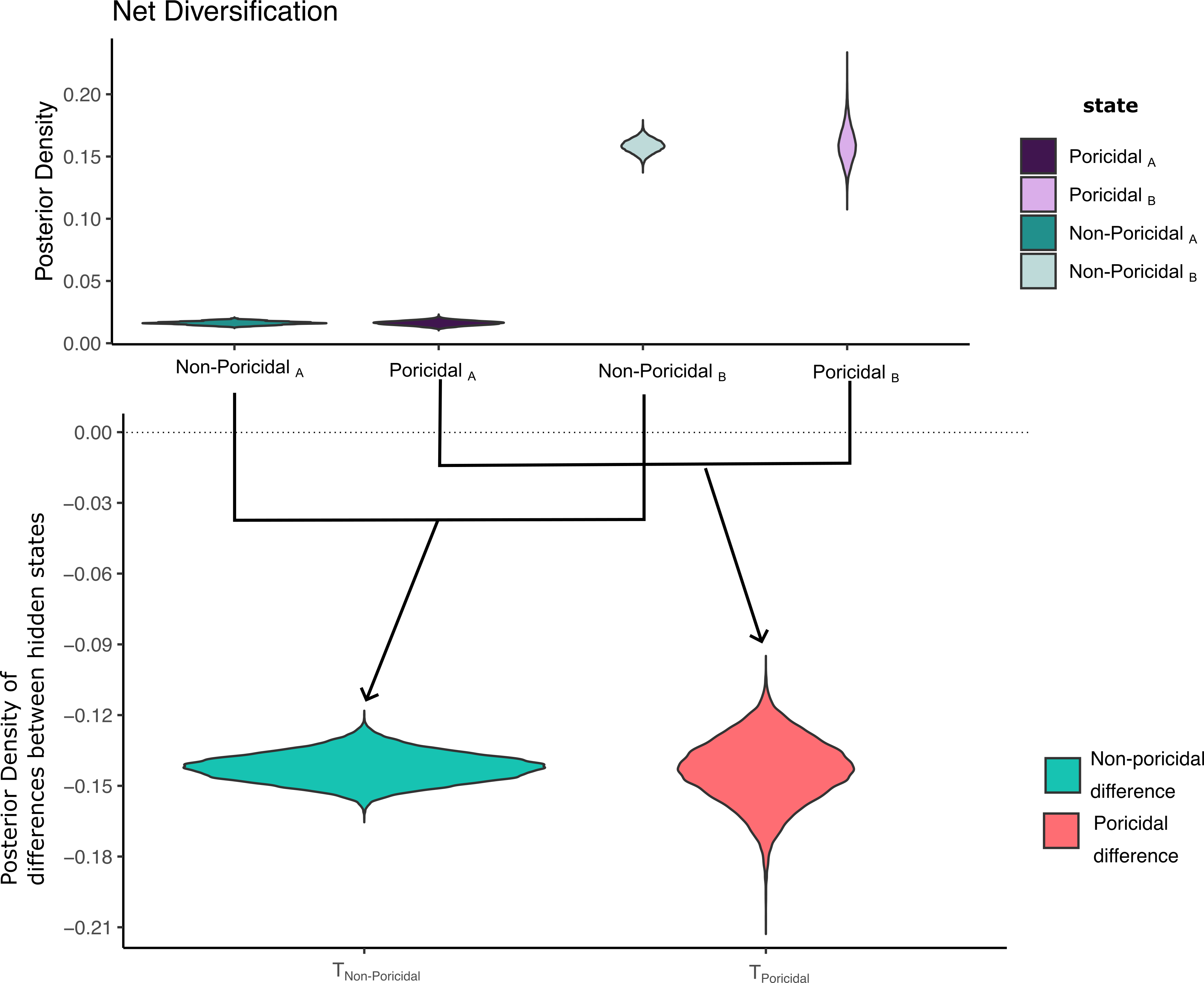

***Figure S3****. Calculating the posterior probability of the test statistics measuring the differences in hidden states A and B. The probability for the test statistics*$T_{Non-poricidal}$*and* $T_{Poricidal}$ *to be zero is negligible, since zero is in the tail of the distribution.*

As we have shown in Figures S2 and S3, building the hypothesis from the CID-2 model in a Bayesian framework only requires transformation of the samples of the posterior distribution. Using these steps, we can then conclude that being poricidal is not linked to diversification at the species-level (see Results in main manuscript). In general, formal testing as we have presented in Figures S2 and S3 is not needed, since posterior distributions allow us to visually compare these estimates from Figures 3 and 6 and infer with high probability differences between states. The main reason why this has not been done in most diversification studies is because the HiSSE R-package (Beaulieu & O’Meara, 2016; Beaulieu *et al.*, 2023) only allows for point estimates (maximum likelihood estimates), which prevent researchers from examining the uncertainty of their fitted parameters. In our case, leveraging the posterior distributions without having to fit more models is key, because the size of the phylogenetic trees presented (10K and 30K tips) makes it computationally complex to fit many models and to calculate the Bayes Information Criterion (BIC). We tried HiSSE (Beaulieu *et al.*, 2023) and castor R-packages (Louca, 2023), both of which failed at convergence of the likelihood due to the numerical difficulty of fitting a state-dependent diversification model in a mega phylogeny. We waited 11 months for convergence of HiSSE’s MCMC in RevBayes, knowing that we would not only get point estimates, but also the full uncertainty of those estimates, and we leverage that uncertainty to compare to the hypothesis of interest as described above.

**Convergence of the MCMC for the HiSSE analysis applied to the species level-tree**

Here we show the convergence of the two runs of the MCMC that we have presented in the main paper for the species-level analysis done with the Janssens et al. (2020) phylogenetic tree, using a sample of 29,246 taxa. Convergence of any MCMC in a large phylogenetic tree is extremely slow and difficult (Fig. S5 and S6) due to the numerical challenges when calculating the very small likelihoods probabilities derived from thousands of branches (i.e., in our tree with 29K tips we have 59,998 branches in which transition probabilities are log transformed and added, resulting in a large log likelihood of –3.7x105, which in its original scale is effectively zero and hard to optimize). Therefore, we waited until we reached an effective sample size (ESS) of more than 200 for only the posterior distribution of the HiSSE model, an 11-month wait, acknowledging that we found many local maxima for the posterior distribution. Many close local maxima were present in the posterior distribution of speciation rate for the non-poricidal A state. All these local maxima have the same consistent result regarding differences in diversification (Fig. S7).

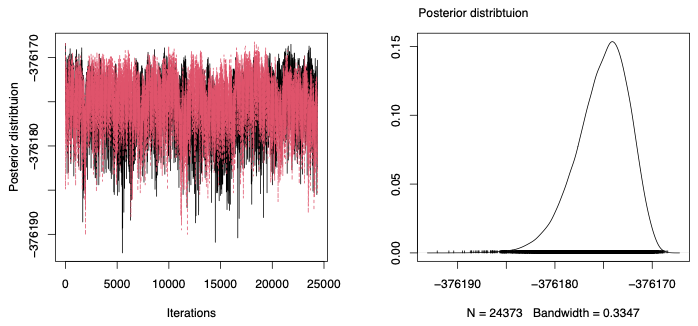

***Figure S4****. Trace plot of the posterior distribution for the HiSSE analyses after discarding the burn-in (burn-in required an 11 month wait).*

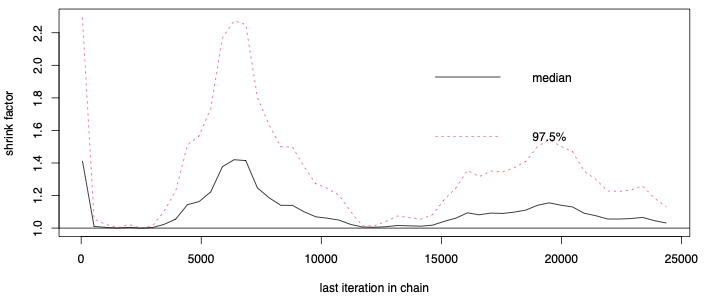

***Figure S5****. Gelman and Rubin diagnostic plot for convergence of the posterior. Convergence of the MCMC is slow given the size of the phylogenetic tree presented. To ensure even sampling, we took samples from the iterations 13000-2500 with a shrink factor below 1.1.*

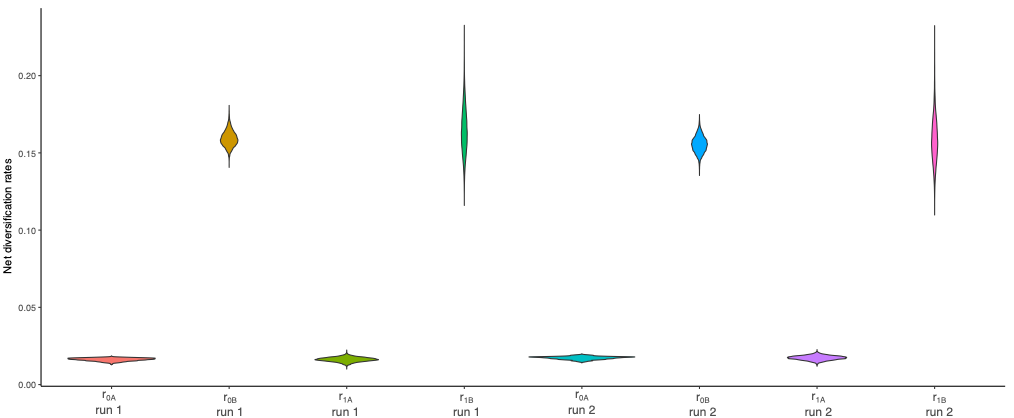

***Figure S6****. Posterior distributions for net diversifications for runs 1 and 2 of the MCMC. As observed above, net diversifications for each of the states are the same in the two runs for the samples obtained. The inference drawn for the two samples is that there are no net diversification differences between state non-poricidal (0) and poricidal (1) within each hidden state. However, differences between A and B exist for each of the main states.*
